# Discrete Transcriptional States Define Biphasic Immune Response and Dynamic CMS Transitions in Colorectal Cancer

**DOI:** 10.64898/2026.02.03.703597

**Authors:** Khalid Ishani, Dechen Wangmo, Atef Ali, Travis Gates, Zejie Yan, Ava P. Gustafson, Ella Boytim, Katie Storey, Paolo Goffredo, Justin Hwang, Subbaya Subramanian

**Author notes:** Corresponding authors **Subbaya Subramanian** University of Minnesota Medical School, Department of Surgery, 11-212 Moos Tower, Mayo Mail Code 195, 420 Delaware Street SE, Minneapolis, MN 55455., **Justin H. Hwang** Masonic Cancer Center, University of Minnesota, Minneapolis, MN. Dwan Building 324 425 E River Parkway Minneapolis, MN 55455.

## Abstract

**Background:** Sequential alterations in *APC*, *KRAS*, *TP53*, and *SMAD4* have been proposed as a framework for colorectal cancer progression. Human colorectal cancer datasets have not revealed the biological transitions associated with these mutations. When examining a cohort of TCGA-colorectal tumors grouped as *AK (APC/KRAS), AKP (APC/KRAS/TP53),* and *AKPS (APC/KRAS/TP53/SMAD4)*, we observed no significant differences in immune-cell composition, four previously defined Consensus Molecular Subtypes (CMS1/2/3/4), or transcriptomic clustering between these genomic groups. Therefore, these canonical alterations do not sufficiently characterize the known properties of metastatic progression in human colorectal cancer.

**Methods:** To overcome these limitations, we developed a genetically defined, organoid-based, orthotopic mouse model whereby mouse colon organoids modeling sequential *APC*, *KRAS*, *TP53*, and *SMAD4* alterations were orthotopically injected into the colon. This was followed by RNA-sequence processing, normalization with DESeq2, differential expression, pathway enrichment, and immune/stromal inference. Gene co-expression modules were identified from variance-stabilized mouse expression data, mapped to 1:1 human orthologs, and summarized as eigengenes. A multinomial logistic regression model trained on mouse eigengenes was applied to TCGA-COAD human tumors to assign them to mouse-informed transcriptomic states (AK-like, AKP-like, AKPS-like), which were then used for downstream visualization and comparative analyses.

**Results:** Whole-transcriptome analysis revealed discrete transcriptional states and immune-cell differences between the organoid *AK/AKP/AKPS* groups. Early *TP53* loss led to strong activation of immune pathways, accompanied by increased infiltration of NK and T cells. As tumors progressed with *SMAD4* loss and metastasis, this immune activity collapsed, giving rise to broad immune suppression. CMS classifications also shifted, with AK tumors resembling epithelial CMS2, AKP tumors displaying immune-rich CMS1 features, and AKPS and metastatic lesions adopting mesenchymal CMS4 characteristics. We then applied a progression-based transcriptomic classifier to 460 human colorectal tumors. This reclassification revealed conserved immune remodeling, CMS transitions, pathway-level differences, and significant differences in patient survival.

**Conclusion:** We show that organoid-derived progression profiles reveal hidden evolutionary structure in human colorectal cancer and provide a transcriptional framework for interpreting metastatic potential and clinical outcomes.

## BACKGROUND

Colorectal cancer (CRC) remains one of the leading causes of cancer-related mortality worldwide, accounting for approximately 10% of all cancer deaths (1). Despite significant advances in early detection, surgical techniques, and adjuvant therapies, the five-year survival rate for patients with metastatic disease remains below 15% (2). A persistent challenge in improving outcomes for CRC patients lies in understanding how tumors evolve from localized lesions to metastatic disease, particularly when accounting for intracellular genomic aberrations and dynamic changes in their immune landscapes and stromal interactions. There is a critical need to define the temporal and mechanistic sequence of transcriptomic, immunologic, and phenotypic transformations using tumor progression models that account for intercellular interactions.

Recent efforts to classify CRC into biologically meaningful subtypes have led to the development of the Consensus Molecular Subtypes (CMS) framework, which captures major transcriptomic and microenvironmental differences based on human tumors. The four CMS categories, CMS1 (immune), CMS2 (canonical), CMS3 (metabolic), and CMS4 (mesenchymal), not only represent distinct biological pathways but also are associated with differences in prognosis, therapeutic responsiveness, and likely mechanisms of resistance (3). For example, CMS1 tumors, which are microsatellite instable (MSI) and immune-rich, are generally more responsive to immunotherapies, whereas CMS4 tumors, characterized by mesenchymal features and stromal infiltration, are associated with poor prognosis and resistance to standard treatments (4). While prognostic, CMS subtypes are only associated with disease stage (3), and it remains unclear whether these subtypes are static or evolve dynamically during tumor progression. Additionally, it is unknown whether specific somatic alterations regulate these phenotypic states or if they are an adaptation to the tumor microenvironment (TME).

Parallel to this growing appreciation of subtype complexity is an expanding body of research recognizing the immune system’s dual role in both suppressing and shaping tumor progression (5). CRC has long been considered an immune cold tumor outside the MSI-high subset (6–10). However, emerging evidence suggests that early stages of colorectal tumors are characterized by more immune activity than previously thought (11). Over time, the TME shifts, and immunosuppressive cell populations play a prominent role in more advanced tumors (12–14). The factors driving these transitions and the timing of their occurrence are still poorly understood. Understanding these patterns is essential for optimizing immunotherapeutic strategies and identifying intervention windows.

A significant obstacle in mapping these dynamic processes in human tumors is the biological and technical complexity of clinical samples. Human CRC samples often consist of heterogeneous cell populations, are influenced by variable host factors, and reflect heterogeneous stages of disease by the time they are sampled (15). Additionally, the immune microenvironment in human tumors is shaped by a lifetime of environmental exposures, microbiome interactions, and systemic immunologic influences, making it challenging to distinguish tumor-associated changes from extrinsic confounders (16–21). Therefore, from genomic data captured at one point in time, it is difficult to discern how alterations in *APC*, *KRAS*, *TP53*, and *SMAD4* modulate tumor–immune interactions and drive state transitions in CRC.

To address this and gain insight into the shifting transcriptomic landscape of CRC as it acquires sequential mutations, we deployed a stepwise organoid-based mouse model of CRC progression. By sequentially introducing driver gene mutations into normal colonic epithelial cells, we established a platform that mimics the genetic and biological trajectory of tumor evolution, from adenomatous lesions (AK) to intermediate carcinomas (AKP) to invasive primary tumors (AKPS) and eventually to distant metastases. This system enables the longitudinal interrogation of molecular and immunologic events associated with CRC progression in a clean, interpretable, and reproducible manner. Guided by these experimental insights, we then reanalyzed human CRC datasets using gene modules derived from transcriptomic data from AK, AKP, and AKPS mouse tumors. This mouse-model-informed classifier successfully mapped each patient tumor to its corresponding mouse-derived state (AK-like, AKP-like, or AKPS-like). This altogether yielded a framework that provides a way to annotate human CRCs and offers a conceptual foundation for identifying critical windows in which immune-based interventions may achieve maximal therapeutic impact or influence patient outcomes.

## METHODS

### Human transcriptomic data acquisition and processing (TCGA-COAD)

Raw gene-level RNA-sequencing counts for primary colorectal adenocarcinoma samples were downloaded from The Cancer Genome Atlas Colorectal Adenocarcinoma (TCGA-COAD) cohort using the Genomic Data Commons (GDC) Application Programming Interface, a publicly accessible NCI-affiliated platform that provides access to genomic data (22). Gene-level counts were converted to transcripts per million (TPM) using GENCODE-annotated gene lengths (23). TPM values were then log₂-transformed. When multiple sequencing aliquots were available for a single patient, barcode strings were parsed, and the sample with the earliest lexicographical plate identifier was retained to ensure that one sample per patient was maintained.

Samples were assigned to the adenomatous lesions (AK), intermediate carcinomas (AKP), or invasive primary tumors (AKPS) categories based on somatic alteration annotations from the GDC (22). TPM values were further converted to fragments per kilobase per million mapped reads (FPKM) and standardized to z-scores. Uniform Manifold Approximation and Projection (UMAP) embeddings were generated from the z-score matrix to visualize the global transcriptomic structure (24).

Immune cell composition was inferred using CIBERSORT with non-log-transformed TPM and HGNC gene symbols as input (25). CMS assignments were obtained using a validated R implementation of the CMS classifier, with log₂(TPM+1) values and human Entrez Gene identifiers supplied as input (3).

### Mouse organoid culture

We maintained normal colon crypts harvested from C57BL/6 WT mice and sh*Apc* organoids as previously described (26,27). Briefly, *AK* (sh*Apc* and *Kras^G12D^*) organoids were generated and provided by Dr. Khazaie. AKP (sh*Apc*, *Kras^G12D^*, and *Trp53 ^KO^*) organoids were generated by introducing an additional *p53* knockout mutation in the AK organoids. The AKPS (sh*Apc*, *Kras ^G12D^*, *Trp53 ^KO^*, and *Smad4^KO^*) organoids, including an additional Smad4 knockout, were from Dr. Westcott (26,27). Normal colon and AK organoids were cultured in complete medium. The AKP organoids were cultured in the same complete media as normal colon organoids with the addition of 13μM of Nutlin-3a. The AKPS organoids were cultured in minimal media (see supplemental methods).

### Colonoscopy-guided organoid injection

Colonoscopy-guided orthotopic injection of organoids was performed, as previously described (28). Briefly, organoids were dissociated into single cells two days before injection and replated. On the day of the injection, organoids were washed in Phosphate Buffered Solution (PBS) and incubated in Dispase (Stem Cell) to dissolve the Matrigel for 10 mins at 37°C. Organoids were washed in PBS and passed through a strainer to ensure that no big chunks of organoids would block the needle. Organoids were resuspended in 10% Matrigel and 90% complete media. Mice were anesthetized using isoflurane (4% for induction and 2% for maintenance). Using the Mainz Coloview mini-endoscopic system (Karl Storz Endoscope), 5000 organoids / 50 μl were injected per mouse through the channel of the colonoscope and inserted into the colonic mucosa at around 30° relative to the colon wall. Successful injection of organoids was confirmed by visualizing a small bubble form within the mucosa.

### Data Processing

Sequencing reads were trimmed for low-quality bases and adapter sequences using Trimmomatic. Processed reads were aligned to the human reference genome (GRCh38, Ensembl release 110) using HISAT2, and resulting alignments were sorted with SAMtools. Raw gene-level read counts were generated using featureCounts and imported as text files for downstream analysis.

### Mouse RNA-sequencing processing and normalization

Raw sequencing counts from the murine tumors were imported as text files. Then, a biomaRt connection to the Ensembl Genes database was established to map the original Ensembl gene IDs to Mouse Gene Identifier (MGI) gene symbols (29). Samples were assembled into a DESeq data frame, and technical replicates were collapsed using DESeq2 (30). The variance stabilizing transformation (VST) was applied to produce a homoscedastic gene-expression matrix (24,25,29). UMAP embeddings were generated from the VST-normalized matrix to visualize genotype-dependent clustering. Sample-to-sample similarity was quantified using Spearman correlation, selected for its robustness to nonlinearity and outliers. The resulting correlation matrix was hierarchically clustered using Euclidean distance and complete linkage. Heatmaps were visualized using the *corrplot* R package (31).

Differential expression analyses were performed on raw counts using DESeq2, with post-hoc effect-size moderation using *lfcShrink* to obtain stable log₂ fold-change estimates (30). Pathway-level differences between genotypes were evaluated using preranked Gene Set Enrichment Analysis (GSEA) with MSigDB Hallmark gene sets (32,33). Immune and stromal cell compositions were inferred from TPM-normalized mouse expression matrices using the immunedeconvframework (34) and the murine MCP-counter method (35). CMS classifications for murine tumors were generated using the MmCMS algorithm (36).

### Cross-species derivation of mouse-informed transcriptomic states in human tumors

#### Data normalization and setup (mouse)

Raw counts of the mouse samples (AK, AKP, AKPS) were first normalized using DESeq2 and transformed with VST to generate a gene × sample expression matrix (30). The VST-normalized mouse dataset served as the basis for the discovery of conserved transcriptional programs

#### Gene co-expression module identification

To capture biologically meaningful variation, the 8,000 most variable genes were selected. Pairwise Spearman correlations were computed between the genes, converted to a distance metric (1 – ρ), and subjected to hierarchical clustering using Ward’s minimum variance method (37). Modules of genes were retained if they contained ≥60 genes and exhibited a minimum intra-module correlation >0.6. These thresholds were chosen based on previous work that used them to identify robust colorectal tumor transcriptional programs (38).

#### Cross-species orthology mapping (mouse to human)

Genes within each murine co-expression module were mapped to 1:1 human orthologs using a validated orthology-mapping resource (23). Modules with <50% successful ortholog recovery were removed from downstream analyses, yielding a high-confidence set of human-interpretable gene modules.

#### Module eigengene calculation

For each retained module, eigengenes were computed as the first principal component of the expression matrix using base R. Eigengenes provided sample-level quantitative summaries of module activity in both mouse and human datasets.

#### Cross-species classification of human TCGA tumors

A multinomial logistic regression model was trained on the mouse eigengene matrix with genotype (AK, AKP, AKPS) as the categorical outcome. The fitted model was applied to eigengenes derived from TCGA-COAD samples to assign each human tumor to its most closely aligned mouse-like transcriptomic class.

#### Downstream analyses using cross-species transcriptomic labels (human)

Human TCGA-COAD samples were re-embedded in UMAP space using z-scored FPKM values, with points colored according to their newly assigned transcriptomic classes. A heatmap of module eigengene expression across human tumor transcriptomic labels was generated using pheatmap (39). A confusion matrix comparing genomic vs. transcriptomic class labels was also visualized using the same framework. Differential gene expression, GSEA, CIBERSORT immune deconvolution, and CMS subtype assignment were repeated as previously described, using the new transcriptomic classifications as the grouping variable for all analyses (24,25,29). All analysis was performed using R version 4.3.3.

## RESULTS

### Human CRC tumors classified by *APC/KRAS/TP53/SMAD4* alterations status show minimal transcriptional or immunologic separation

We first aimed to determine whether canonical driver mutations in CRCs could be used to demarcate biological states in human tumors, thereby annotating the 460 CRC samples from TCGA using mutational groupings: *APC* loss, *KRAS* alterations or amplifications, *TP53* alteration or loss, and *SMAD4* loss. Tumors were genomically stratified as AK (*APC*+*KRAS*), AKP (*APC*+*KRAS*+*TP53*), or AKPS (*APC*+*KRAS*+*TP53*+*SMAD4*). This ultimately captured the subsets of 55 tumors as AK, 73 as AKP, and only 11 as AKPS (**Figure 1A**), underscoring the relative sparsity of human CRCs expressing an AKPS genotype. However, despite clear genomic delineation, transcriptomic profiling failed to reveal biologically meaningful separation between these subgroups. Specifically, we performed dimensionality reduction of the whole transcriptomes from these samples using UMAP, which showed extensive overlap and no distinct clusters by genomic category (**Figure 1B**). This indicated that the *APC/KRAS/TP53/SMAD4* alteration status alone, implicated in paradigms of adenoma-carcinoma progression, is insufficient to demarcate the landscape of human CRC.

**Figure 1.**
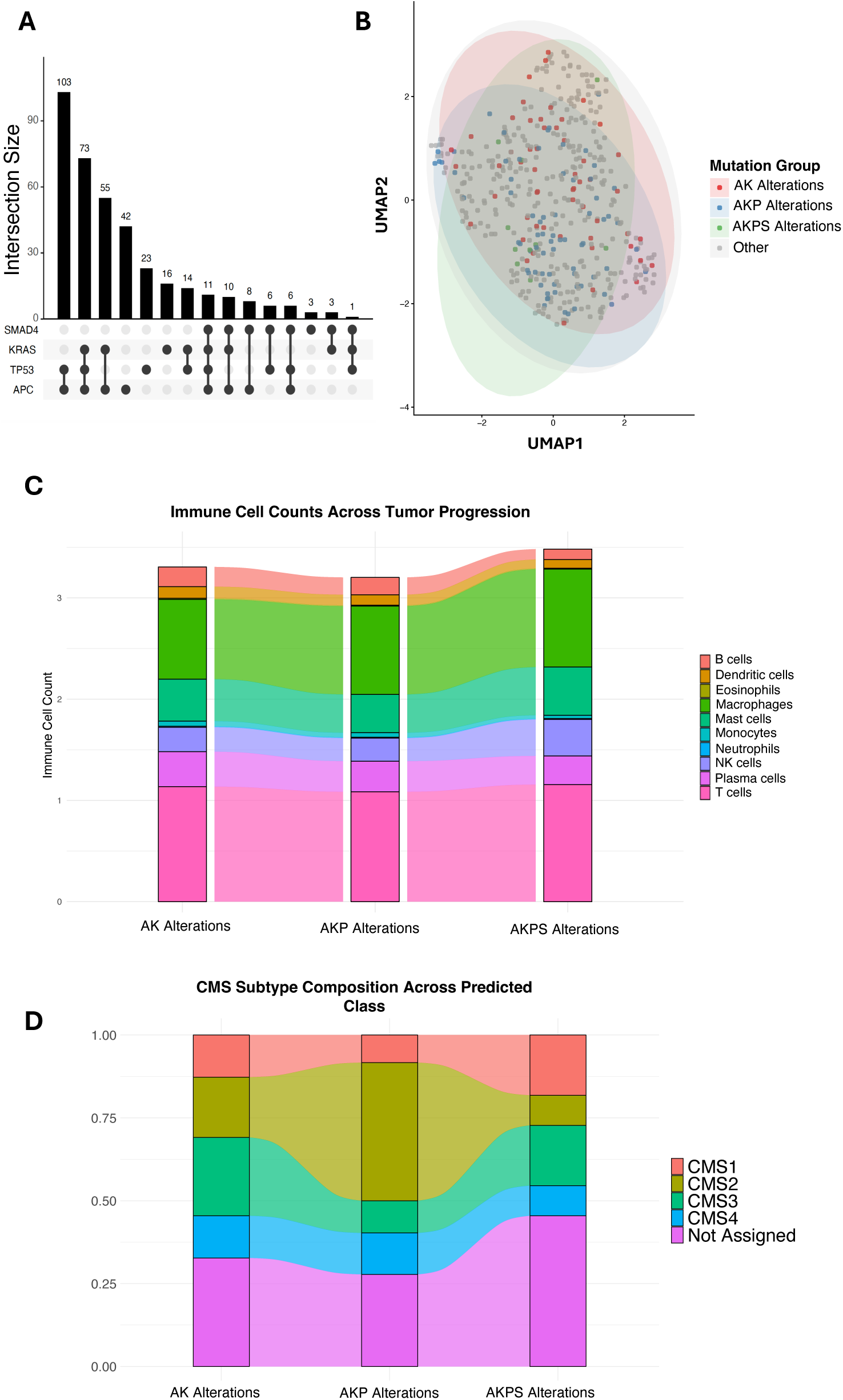
Transcriptomic and Gene Expression Changes Across Human Tumor Stages Labeled by Genomic Alterations. **(A)** UpSet plot showing the number of samples with various combinations of mutations in the genes APC, TP53, KRAS, and SMAD4. A total of 374 tumor samples corresponding to 372 unique patients were analyzed. Two patients contributed more than one biologically distinct tumor specimen (primary versus recurrent or metastatic); no technical replicates were included. One of the samples from the AKP genomic alteration group was unavailable in the R API. Therefore, in all analyses comparing AK alterations, AKP alterations, and AKPS alterations groups, 138 biological replicates were used. **(B)** Uniform Manifold Approximation and Projection (UMAP) score plot showing lack of stage-specific (AK alterations, AKP alterations, AKPS alterations) clustering of transcriptomes along major variance axes in human tumor samples, where the labels are based solely on genomic alteration. The RNA counts were converted to z-scores so each gene had mean=0, standard deviation=1 across samples prior to UMAP implementation. A total of 460 tumor samples corresponding to 458 unique patients were analyzed. Two patients contributed more than one biologically distinct tumor specimen (primary versus recurrent or metastatic); no technical replicates were included. **(C)** Alluvial plot showing relative changes in the proportion of immune cell subtypes across sequential stages of the human model, where labels are based on genomic alteration only. A total of 139 tumor samples corresponding to 138 unique patients were analyzed. No technical replicates were included. **(D)** Alluvial plot showing relative changes in the proportion of Conesus Molecular Subtype (CMS) subtypes across sequential stages of the human model, where labels are based on genomic alteration only. A total of 139 tumor samples corresponding to 138 unique patients were analyzed. No technical replicates were included.

We next evaluated whether immune microenvironment features differed across these genomically defined groups. Immune deconvolution followed by alluvial mapping revealed broad overlap in relative proportions of major immune cell types, including T cells, B cells, macrophages, and neutrophils, across AK, AKP, and AKPS tumors (**Figure 1C**). No consistent or directional shifts in immune composition were observed as tumors accumulated additional driver mutations.

We next assessed whether the AK, AKP, and AKPS genomic states correspond to shifts in established human CRC subtypes. The distribution of CMS types (CMS1–CMS4) remained broadly similar across all three groups, with CMS2 and CMS3 consistently accounting for the most significant fractions (**Figure 1D**). CMS1 and CMS4 subtypes were present at lower and relatively uniform proportions. Although CMS2 showed a significant increase in the AKP state and CMS3 displayed a modest decrease, these changes were difficult to contextualize with the lack of immune environment and transcriptomic differences between the three genomic groups.

### Generation of the AKPS organoid model recapitulates stepwise genetic progression, adenocarcinoma histopathology, and spontaneous metastasis

To establish an experimentally tractable system to model sequential CRC progression, we used a murine organoid model that enabled controlled, stepwise introduction of *Apc*, *Kras*, *Tp53*, and *Smad4* alterations. The conceptual framework of this system mirrors the canonical progression from normal colonic epithelium to invasive CRC, with each genetic alteration representing a discrete biological transition (**Figure 2A-B**).

**Figure 2.**
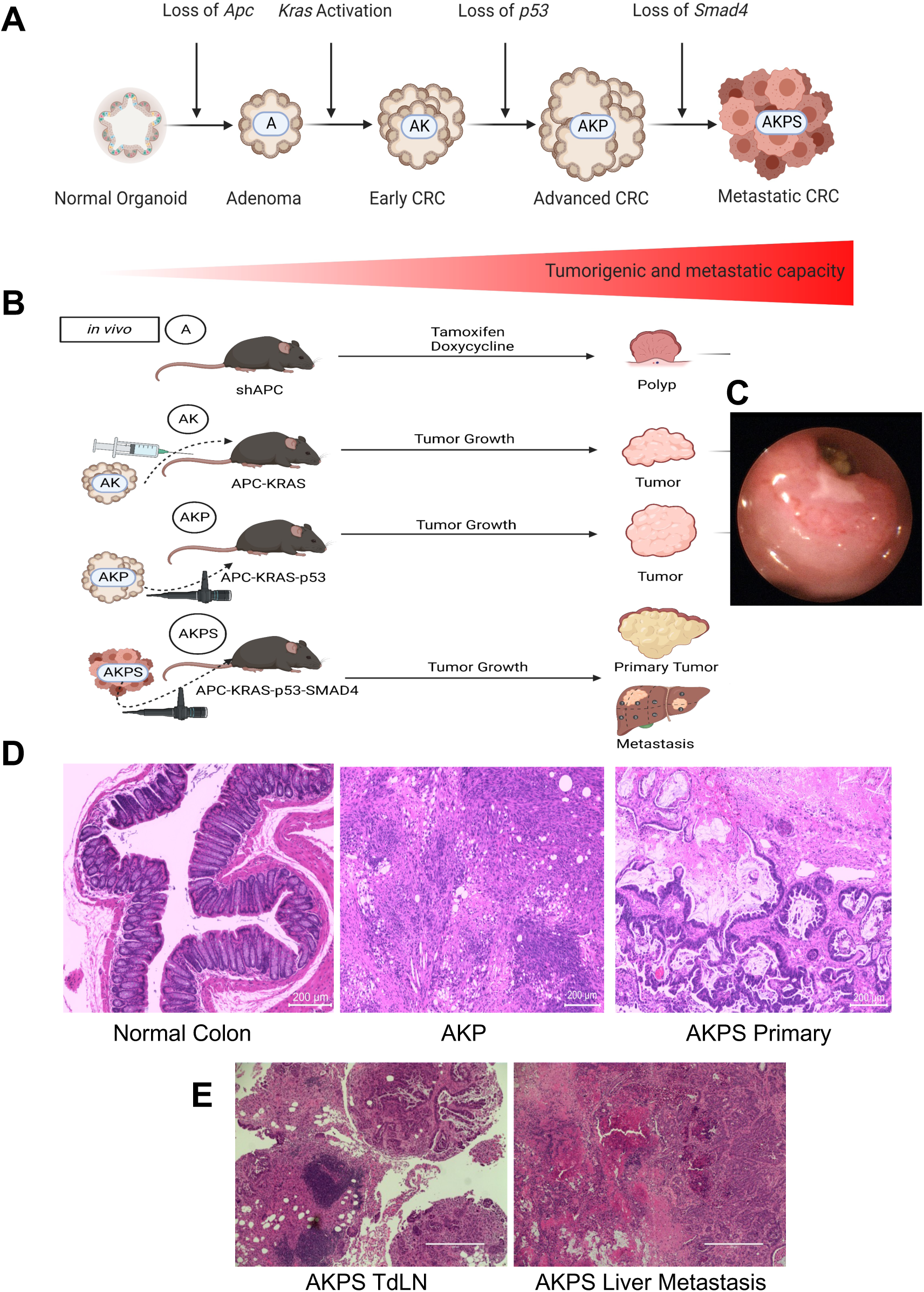
Sequential Genetic and Histologic Progression in the Murine Organoid model of CRC. **(A-B)** Conceptual illustration of the stepwise progression from normal colon epithelium to metastatic CRC, highlighting the sequential acquisition of CRC driver gene mutations. **(C)** Representative endoscopy image of colon tumor. **(D-E)** Representative H&E stained sections showing normal colon, AKP, AKPS primary tumor, AKPS metastasis to a tumor-draining lymph node (TdLN), and AKPS metastasis to the liver.

Colonoscopic visualization of the mouse colon revealed a polypoid intraluminal mass arising from the mucosal surface, consistent with adenomatous and early neoplastic growth (**Figure 2C)**. These lesions exhibited vascularity and mucosal disruption reminiscent of human adenomas and early carcinomas. These features indicate that the engineered mutations collectively drive progression from benign epithelial hyperplasia to high-grade invasive carcinoma, validating the histopathologic fidelity of the model. Importantly, AKPS tumors exhibited spontaneous metastatic potential (27), a key criterion for modeling advanced-stage disease. H&E-stained sections of the mouse colonic tissue showed early lesions with polypoid mucosal expansion and glandular irregularity (**Figure 2D-E)**. We have demonstrated that AKPS organoids serve as the basis for a metastatic CRC model that recapitulates advanced CRC in vivo (27). AKPS primary tumors exhibited invasive adenocarcinoma morphology, including markedly dysplastic epithelium (**Figure 2D-E**).

### Organoid-derived tumors exhibit clear transcriptional stratification across defined progression stages

To determine whether sequential acquisition of *Apc*, *Kras*, *Tp53*, and *Smad4* alterations generates discrete transcriptional states under controlled biological conditions, we performed RNA sequencing of tissues derived from our organoid-based progression model. Uniform Manifold Approximation and Projection (UMAP) analysis of all samples revealed striking stage-specific segregation, in contrast to the minimal separation observed in human tumors grouped solely by genotype (**Figure 3A**). Normal colon (n = 3), polyp tissue (n = 3), AK tumors (n = 3), AKP tumors (n = 3; two biological and one technical replicate), AKPS tumors (n = 3), metastases to tumor-draining lymph nodes (n = 3 technical replicates), distant organ metastases (n = 3 technical replicates), and lung metastases (n = 3 technical replicates) each formed compact, internally consistent clusters. These results indicate that the organoid-based system captures differences aligned with defined genetic and phenotypic states.

**Figure 3.**
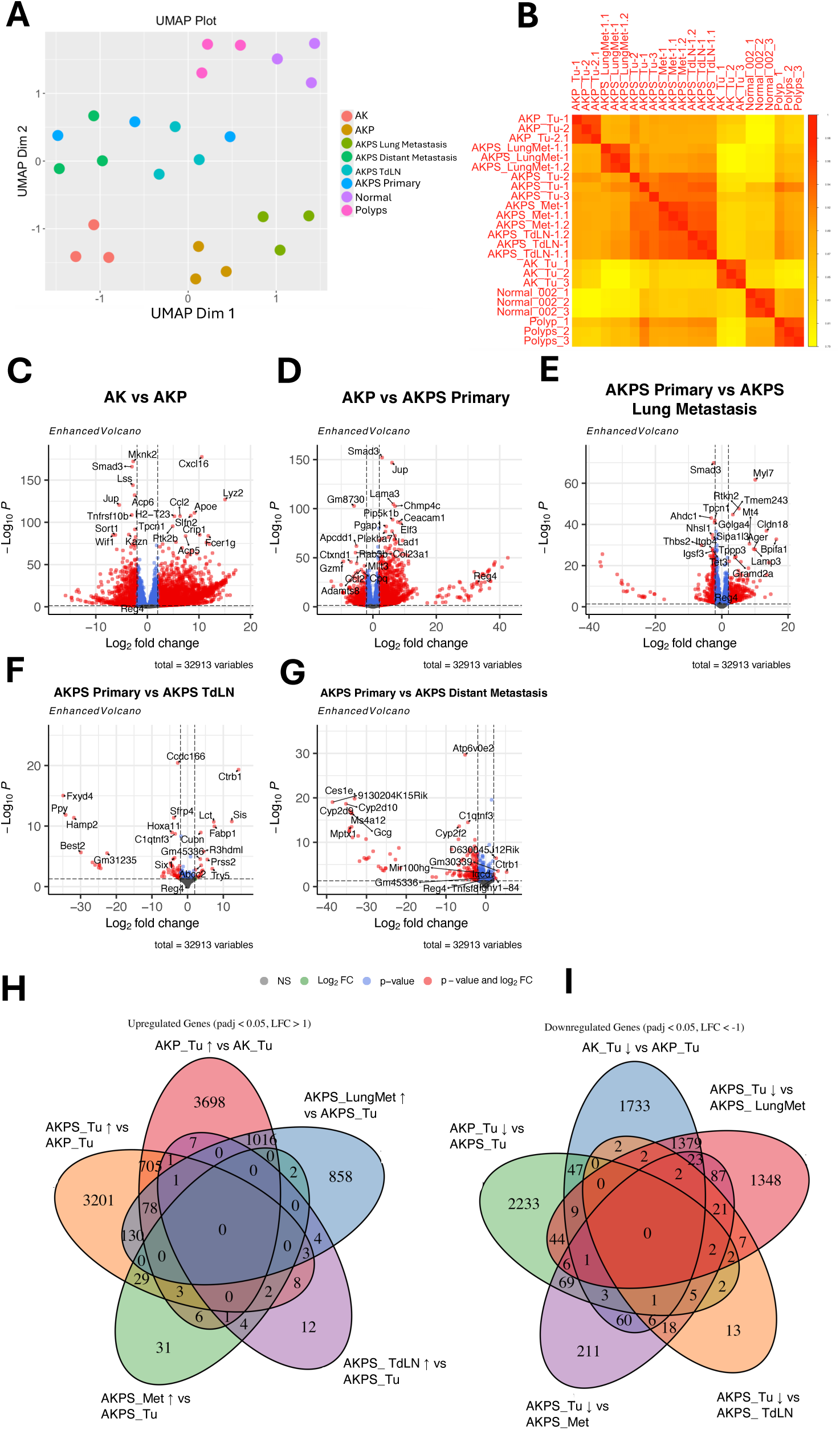
Transcriptomic Relationships and Gene Expression Changes Across Tumor Progression Stages. **(A)** Uniform Manifold Approximation and Projection (UMAP) plot showing stage-specific clustering of transcriptomes along major variance axes. **(B)** Correlation matrix of RNA sequencing data across all progression stages in the organoid model, illustrating high within-stage similarity and distinct between-stage profiles. **(C-G)** Volcano plots depicting differentially expressed genes for each sequential progression step. Genes meeting thresholds for both adjusted *P* value and log2-fold change are shown in red; those significant only for adjusted *P* value in blue; nonsignificant genes in gray. P-values were calculated using the Wald test. **(H-I)** 5-way Ven diagrams summarizing the number of genes statistically significantly (alpha=0.05) upregulated (figure H) and downregulated (figure I) in the various cell tumor progression comparisons. The ‘Mets’ category combines distant metastases and lung metastases. P-values were calculated using the Wald test. **Biological and technical replicates:** Each comparison was performed using biological replicates, with technical replicates present only for selected AKPS metastatic samples. Biological replicate counts were: Normal colon (n=3), Polyps (n=3), AK (n = 3), AKP (n = 2), AKPS primary tumors (n = 3), AKPS distant metastasis (n = 1), AKPS lung metastasis (n = 1), and AKPS TdLN (n = 1). Technical replicates were generated only for AKPS lung metastasis (3 runs), AKPS distant metastasis (3 runs), AKPS_TdLN (3 runs), and AKP_Tu-2 (2 runs). All analyses were performed on biological replicate–level summaries.

A pairwise sample correlation matrix corroborated these findings, demonstrating high intra-stage concordance and pronounced inter-stage divergence (**Figure 3B**). Samples within each category displayed correlation coefficients approaching unity, whereas correlations across categories decreased sharply, consistent with substantial transcriptomic rewiring at each biological transition. This pattern reinforces the notion that the organoid system produces robust and reproducible transcriptional signatures that reflect discrete stages of tumor evolution.

To identify the specific molecular changes underlying these separations, we performed differential gene-expression analysis across all sequential transitions. Volcano plots revealed extensive gene-expression remodeling at each step, with hundreds of transcripts significantly altered in AK vs. AKP tumors, AKP vs. AKPS tumors, and AKPS tumors vs. their metastatic derivatives (**Figures 3C–G**). Both upregulated and downregulated genes were represented, demonstrating that each stage is shaped by dynamic activation and suppression of distinct biological pathways rather than by uniform intensification of a single program. The breadth of transcriptional changes observed across these transitions underscores the profound functional consequences of each added driver alteration.

To integrate these findings and determine the extent to which gene-expression responses were shared or unique across progression steps, we quantified overlapping differentially expressed genes using five-way Venn diagrams. The analysis revealed limited overlap across stages for both upregulated (**Figure 3H**) and downregulated (**Figure 3I**) genes, indicating that most transcriptional alterations were stage restricted. Only a small subset of genes was consistently altered across multiple transitions, suggesting that tumor evolution in this system is not driven by incremental amplification of early alterations but rather by successive activation of distinct transcriptional programs.

### GSEA reveals a biphasic immune and EMT program during CRC progression

To define the coordinated biological programs underlying sequential tumor evolution in the organoid model, we performed Gene Set Enrichment Analysis (GSEA) using the Hallmark pathway collection across all stage-to-stage comparisons. This analysis revealed a biphasic pattern of early activation and later suppression of inflammatory and epithelial–mesenchymal transition (EMT) programs, suggesting that progression from tumor initiation to metastatic dissemination is driven by discrete, temporally ordered transcriptional waves.

The first significant wave emerged during the transition from AK to AKP tumors, which showed robust enrichment of immune and inflammatory pathways. Heatmap visualization of normalized enrichment scores (NES) demonstrated that interferon-γ and interferon-α responses, IL2-STAT5 and IL6-JAK-STAT3 signaling, complement activation, allograft rejection, and generalized inflammatory response signatures were all significantly upregulated in AKP relative to AK tumors (**Figure 4A**). The breadth and magnitude of these immune-related enrichments indicate that introduction of *Tp53* loss in the context of *Apc* and *Kras* alterations triggers a coordinated inflammatory transcriptional program that marks a critical inflection point in tumor evolution. This immune activation was accompanied by upregulation of the EMT Hallmark gene set during the same transition (**Figure 4B**), suggesting that the AK-to-AKP step involves both heightened inflammatory signaling and acquisition of mesenchymal-like features.

**Figure 4.**
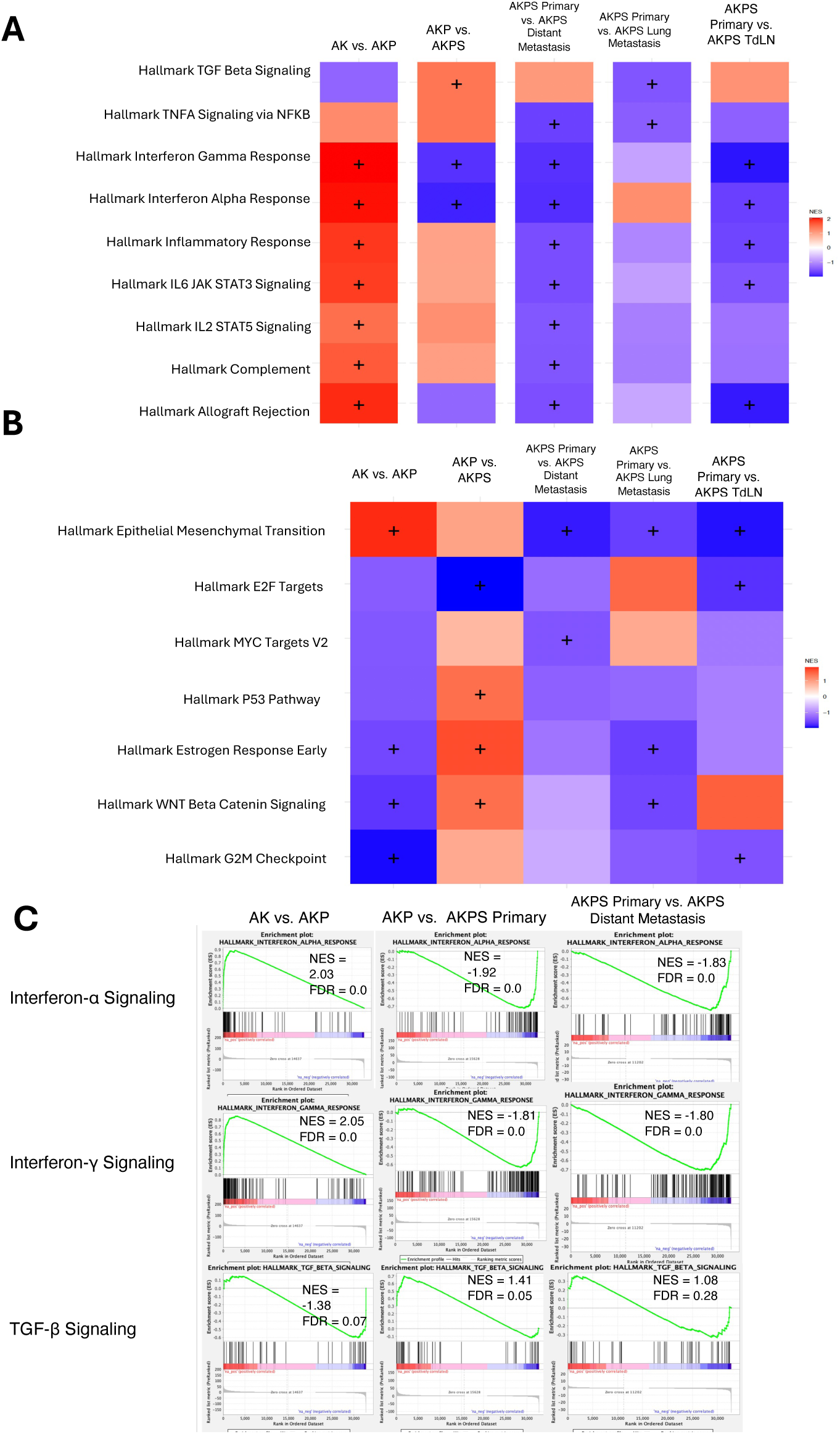
Hallmark Pathway Dynamics Across Sequential Tumor Progression. **(A)** Heat map of normalized enrichment scores (NES) for selected inflammatory Hallmark pathways identified by pre-ranked Gene Set Enrichment Analysis (GSEA) across stage-to-stage comparisons in the organoid model. A positive NES indicates enrichment in the later stage of each comparison. Plus signs denote pathways reaching statistical significance (FDR-adjusted *q* < 0.05). Significance was assessed using a permutation-based weighted Kolmogorov–Smirnov–like enrichment statistic with gene set permutations, with false discovery rate (FDR) correction applied across all tested pathways. **(B)** Heat map of NES for selected oncogenic Hallmark pathways identified by pre-ranked GSEA across the same stage-to-stage comparisons. A positive NES indicates enrichment in the later stage. Plus signs denote statistically significant pathways (FDR-adjusted *q* < 0.05), with significance assessed using the same permutation-based weighted enrichment statistic and FDR correction as in (A). **(C)** Grid of representative GSEA enrichment plots. Comparisons are shown by column: AK vs. AKP, AKP vs. AKPS, and AKPS vs. distant metastasis. Pathways are organized by row: interferon-α signaling, interferon-γ signaling, and TGF-β signaling. Enrichment significance was determined using pre-ranked GSEA with gene set permutations and FDR correction across all gene sets. **Biological and technical replicates:** Biological replicates included AK (n = 3), AKP (n = 2), AKPS primary tumors (n = 3), AKPS distant metastasis (n = 1), AKPS lung metastasis (n = 1), and AKPS tumor-draining lymph node (TdLN; n = 1). Technical replicates were generated only for AKPS lung metastasis, AKPS distant metastasis, AKPS TdLN (each 3 runs), and AKP_Tu-2 (2 runs). All analyses were performed on biological replicate–level summaries.

In contrast, a second and opposing wave was observed during progression from AKPS primary tumors to metastatic lesions. All comparisons between AKPS tumors and their metastatic derivatives, including distant organ metastases, lung metastases, and tumor-draining lymph node metastases, showed significant downregulation of inflammatory pathways activated earlier in progression (**Figure 4A**). The directionality of these changes, reflected in strongly negative NES values, suggests that metastatic dissemination is associated with broad suppression of interferon signaling and other immune response programs. A parallel pattern was observed for EMT pathway activity. Although EMT was activated earlier during tumor progression, all metastatic comparisons showed significant downregulation of the EMT Hallmark signature relative to AKPS primary tumors (**Figure 4B**).

Enrichment plots for representative pathways, including interferon-α signaling, interferon-γ signaling, and TGFβ signaling, further illustrate this biphasic pattern, with marked pathway activation during the AK-to-AKP transition, stabilization or modest change during the AKP-to-AKPS transition, and pronounced suppression in metastatic comparisons (**Figure 4C**).

### Immune cell abundance and subtype composition shift in a stage-specific manner

To determine how the immune microenvironment evolves alongside defined genetic and phenotypic transitions during organoid progression, we performed immune cell deconvolution using transcriptome-derived MCP-counter estimates. This analysis revealed a dynamic, non-linear pattern of immune engagement throughout tumor development. A heatmap of relative immune cell densities showed a pronounced increase in total immune infiltration during the transition from AK to AKP tumors, with the most substantial contributions from NK cells and T-cell subsets (**Figure 5A**). This increase was not restricted to a single lineage; instead, it reflected coordinated recruitment or expansion of cytotoxic T cells and NK cells, as confirmed by alluvial mapping of immune subtype counts (**Figure 5B**). These data indicate that Tp53 loss in the *Apc*–*Kras* background is associated with a significant and broad influx of immune effectors.

**Figure 5.**
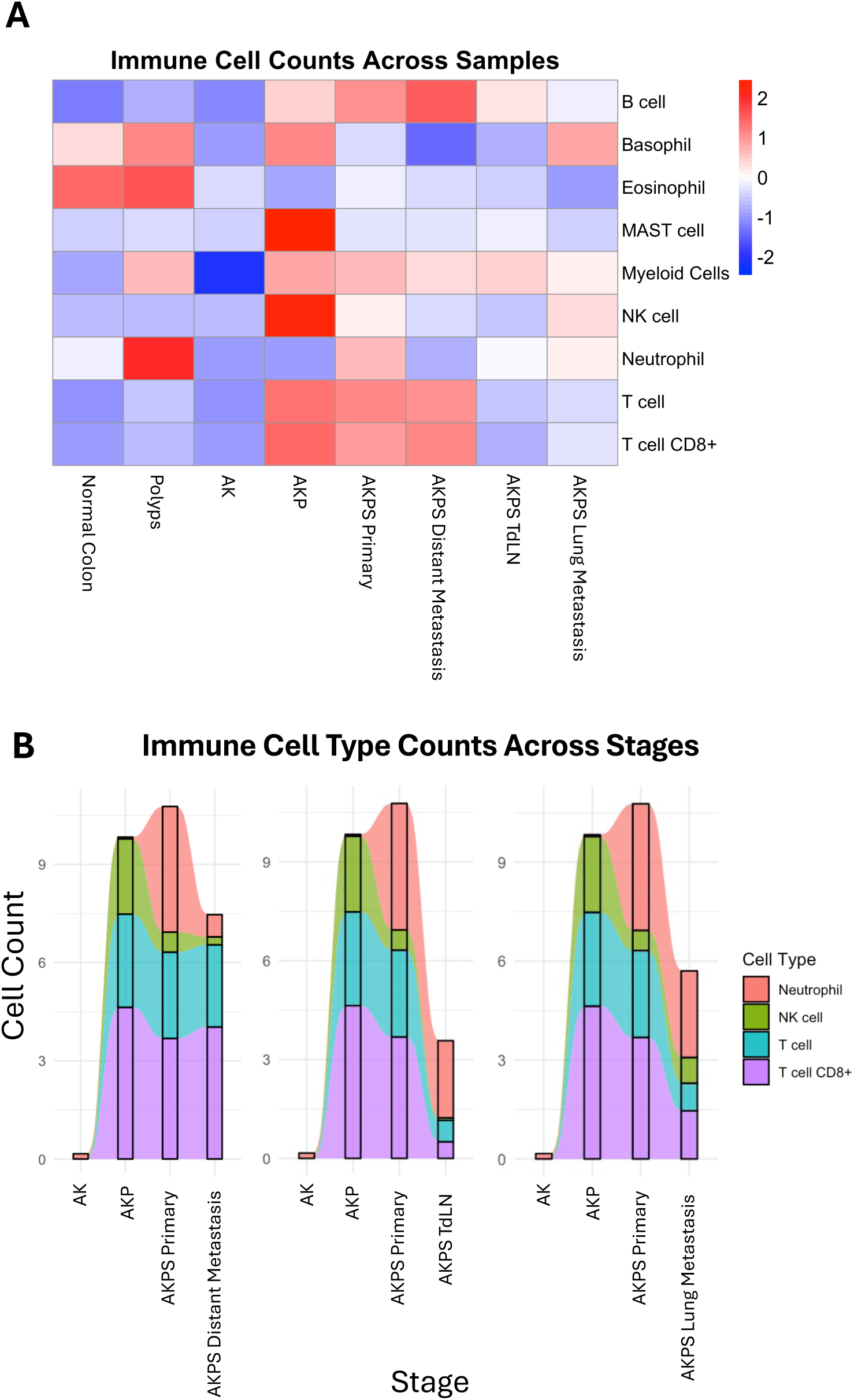
Evolution of Immune Cell Composition Across Tumor Progression Stages. **(A)** Heat map of relative immune cell density. Positive values indicate higher density compared with the preceding stage. Immune cell estimates were derived from RNA sequencing data using the MCP-counter R package. **(B)** Alluvial plot showing relative changes in the count of immune cell subtypes across sequential stages of the organoid model. **Biological and technical replicates:** Each comparison was performed using biological replicates, with technical replicates present only for selected AKPS metastatic samples. Biological replicate counts were: Normal colon (n=3), Polyps (n=3), AK (n = 3), AKP (n = 2), AKPS primary tumors (n = 3), AKPS distant metastasis (n = 1), AKPS lung metastasis (n = 1), and AKPS TdLN (n = 1). Technical replicates were generated only for AKPS lung metastasis (3 runs), AKPS distant metastasis (3 runs), AKPS_TdLN (3 runs), and AKP_Tu-2 (2 runs). For visualization purposes only, for both the alluvial plots and the heatmap, technical and biological replicates were averaged together.

In contrast, the shift from AKP to AKPS tumors did not substantially alter the overall magnitude of immune infiltration but did produce measurable alterations in immune composition. During this interval, T-cell abundance declined while neutrophil representation increased (**Figure 5A-B)**, suggesting a remodeling of the immune environment toward a more inflammatory yet potentially less effective antitumor profile. This compositional shift, despite stable overall immune density, points to qualitative rather than quantitative changes in the immune microenvironment as tumors lose SMAD4 function.

A third distinct pattern emerged during metastatic dissemination. All comparisons involving AKPS primary tumors and their metastatic derivatives, including distant organ metastases, tumor-draining lymph node metastases, and lung metastases, showed a consistent and global reduction in immune infiltration (**Figure 5A–B**). The decrease affected nearly all major immune lineages, indicating the establishment of an immune-suppressed microenvironment within metastatic niches. The convergence of reduced immune abundance across multiple anatomical sites suggests that immune evasion or exclusion is a hallmark of the metastatic state in our model.

### CMS subtype assignments track with transcriptional and immune remodeling in the organoid model

To determine whether established CRC transcriptional subtypes emerge in parallel with the molecular and immunologic transitions observed in the organoid model, we applied the mouse-adapted CMS types (MmCMS) classifier to all samples. This analysis revealed a coordinated and stage-specific progression through CMS categories that aligned with the underlying transcriptional and immune landscapes (**Figure 6**). Tumors harboring *Apc* and *Kras* alterations alone consistently mapped to CMS2, a subtype characterized by canonical epithelial programs and minimal immune activation. Introduction of *TP53* loss shifted classification to CMS1, the immune-rich subtype, mirroring the substantial immune influx and broad interferon pathway activation observed during this transition.

**Figure 6.**
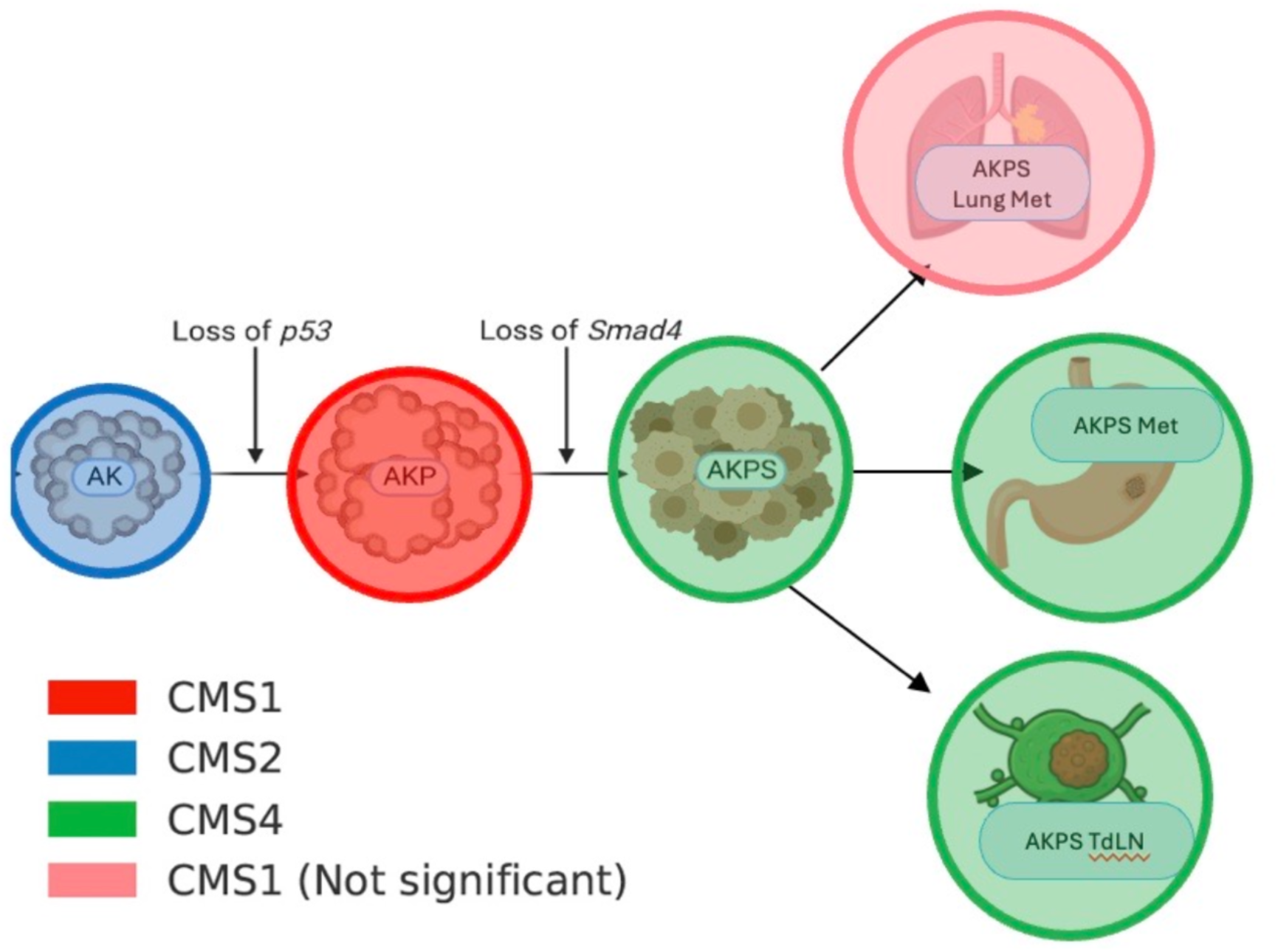
Consensus Molecular Subtype Assignments Across Tumor Progression Stages. Qualitative flow diagram of the predicted Consensus Molecular Subtype (CMS) classifications for individual tumor samples at each stage of the organoid model, determined using the MmCMS R package. Statistical significance was determined using a threshold of alpha = 0.05 after adjustment for the false discovery rate (FDR). N=3 for each sample size, and all tumor samples were classified as the same CMS subtype **(Supplementary Table 1)**.

The addition of *Smad4* loss produced a second major shift: all AKPS primary tumors were classified as CMS4, the mesenchymal and fibroinflammatory subtype. CMS4 assignments coincided with the EMT activation observed during earlier progression.

In contrast, lung metastases displayed an atypical pattern. Although these samples were nominally assigned to CMS1, their assignments carried high false discovery rates and did not reach significance, indicating weak or ambiguous subtype identity. Collectively, these data demonstrate that CMS subtype identity is not a fixed property dictated solely by genotype but instead reflects dynamic tumor–immune–stromal interactions that evolve during carcinoma progression (**Figure 6**).

### Cross-species classification reveals that human CRC tumors follow the same AK–AKP–AKPS immune trajectory

Given the robust stage-specific transcriptional and immune transitions observed in the organoid model, we next sought to determine whether human colorectal tumors exhibit analogous progression states. To do so, we applied a cross-species classification framework in which transcriptional modules derived from the organoid dataset were projected onto TCGA colorectal tumors. (38) Each human tumor was assigned to AK-like, AKP-like, AKPS-like, or none based on the expression patterns of conserved module signatures, allowing us to classify tumors according to functional transcriptional states rather than mutation-defined labels.

Module eigengene analysis demonstrated that the six identified transcriptional modules exhibited distinct and nonredundant patterns across human samples. A heatmap of eigengene values showed that module 4 exhibited the strongest discriminative power among the predicted classes, but no single module fully determined class identity (**Figure 7A**). This multidimensional structure indicates that the AK-like, AKP-like, and AKPS-like transcriptomic states arise from the coordinated contributions of several gene-expression programs rather than a single dominant signature. Such distributed module weighting enhances the classifier’s biological robustness and reduces the likelihood that spurious or lineage-restricted genes are driving classification.

**Figure 7.**
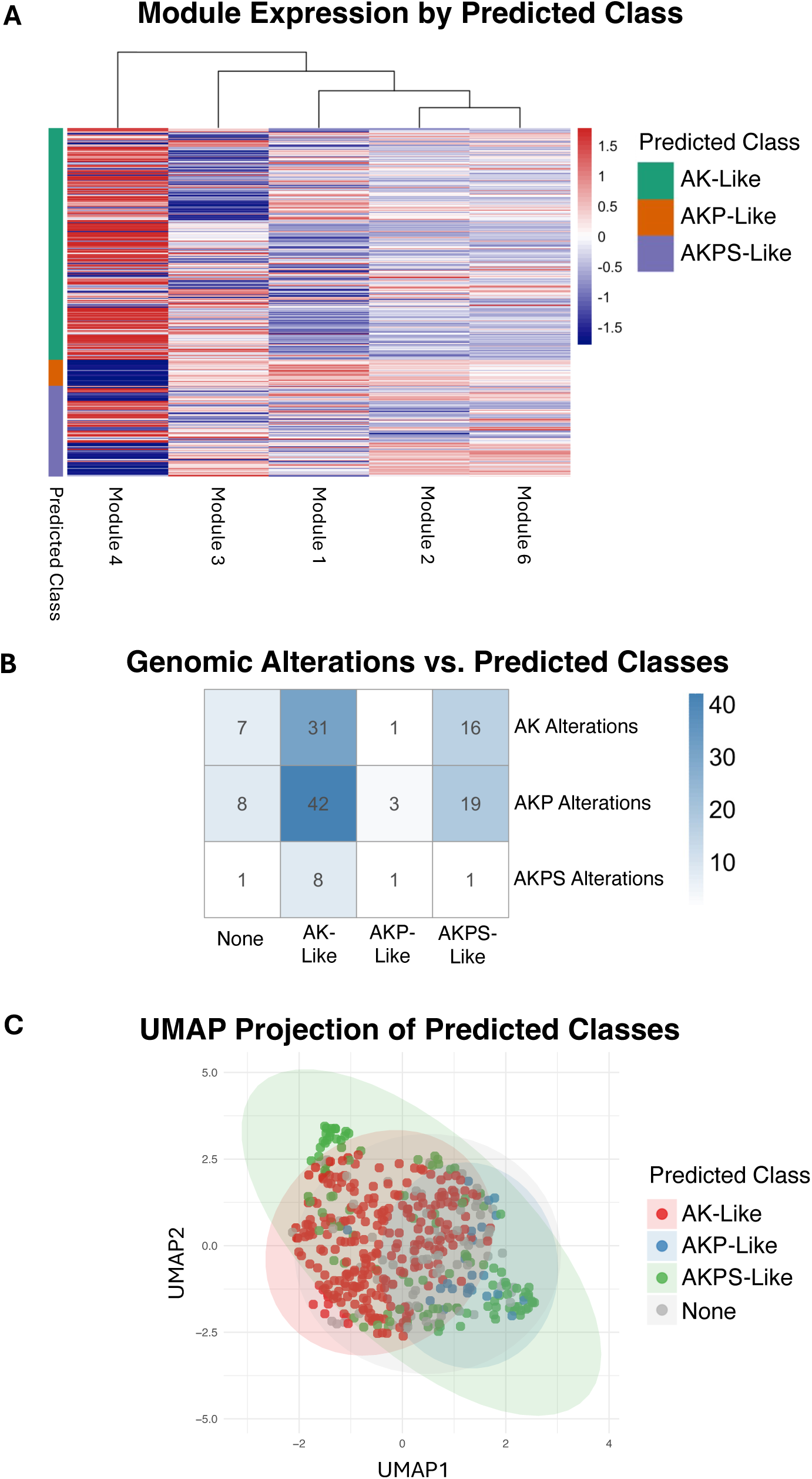
Transcriptomic Clustering of Human Tumor Progression Stages Labeled by Cross-Species Classification. **(A)** Heatmap illustrating the relative expression of modules of genes for each human sample, separated by predicted class based on Cross-Species Classification. **(B)** Confusion matrix illustrating how human samples labeled AK-alterations, AKP-alterations, AKPS-alterations based on genomic classification (y-axis) were classified into predicted classes (x-axis) **(C)** Uniform Manifold Approximation and Projection (UMAP) score plot showing the improved stage-specific clustering of transcriptomes along major variance axes in human tumor samples, where the labels are based on predicted classes (AK-like, AKP-like, AKPS-like). The RNA counts were converted to z-scores, so each gene had a mean of 0 and a standard deviation of 1 across samples prior to UMAP implementation. **Biological and technical replicates:** Each comparison was performed using biological replicates. Biological replicate counts for the various transcriptomic groups were AK-like (n=248), AKP-like (n=22), AKPS-like (n=118), and None (n=72). Of the total 460 tumor samples analyzed, 458 corresponded to unique patients. Two patients contributed more than one biologically distinct tumor specimen (primary versus recurrent or metastatic); no technical replicates were included.

Reclassification of the TCGA tumors using these conserved modules produced a markedly different organizational structure than the initial mutation-based grouping. A confusion matrix comparing genomic labels with predicted transcriptional classes demonstrated widespread discordance between the two approaches (**Figure 7B**). Notably, most tumors, irrespective of their *APC/KRAS/TP53/SMAD4* alteration patterns, mapped to the AKP-like transcriptional state.

Importantly, projecting human tumors into these transcriptomically defined states resolved the lack of structure observed when tumors were stratified by genotype alone. UMAP analysis based on AK-like, AKP-like, and AKPS-like assignments yielded more clearly separated clusters with increased stage-specific organization (**Figure 7C**), in stark contrast to the diffuse, overlapping transcriptomes shown in Figure 1B.

### Human tumors show conserved early immune activation and late immune suppression

Having reassigned TCGA tumors into AK-like, AKP-like, and AKPS-like transcriptional states using cross-species classification, we next tested whether human tumors recapitulate the biphasic immune dynamics identified in the organoid model (**Figure 8A-B)**. GSEA revealed a highly conserved pattern of early immune activation followed by later immune attenuation (**Figure 8C)**. During the transition from AK-like to AKP-like tumors, hallmark inflammatory pathways showed strong, statistically significant enrichment in the AKP-like state. Pathways consistently upregulated included allograft rejection, complement activation, IL2-STAT5 signaling, IL6-JAK-STAT3 signaling, interferon-γ and interferon-α responses, TNFα signaling, and TGFβ signaling (**Figure 8C**). A second wave of immune modulation became apparent when comparing AKP-like to AKPS-like tumors. In contrast to the robust activation observed in the earlier transition, inflammatory pathways were broadly downregulated in AKPS-like tumors, except for complement activation, IL-2 signaling, and TNFα, which were upregulated (**Figure 8C**).

**Figure 8.**
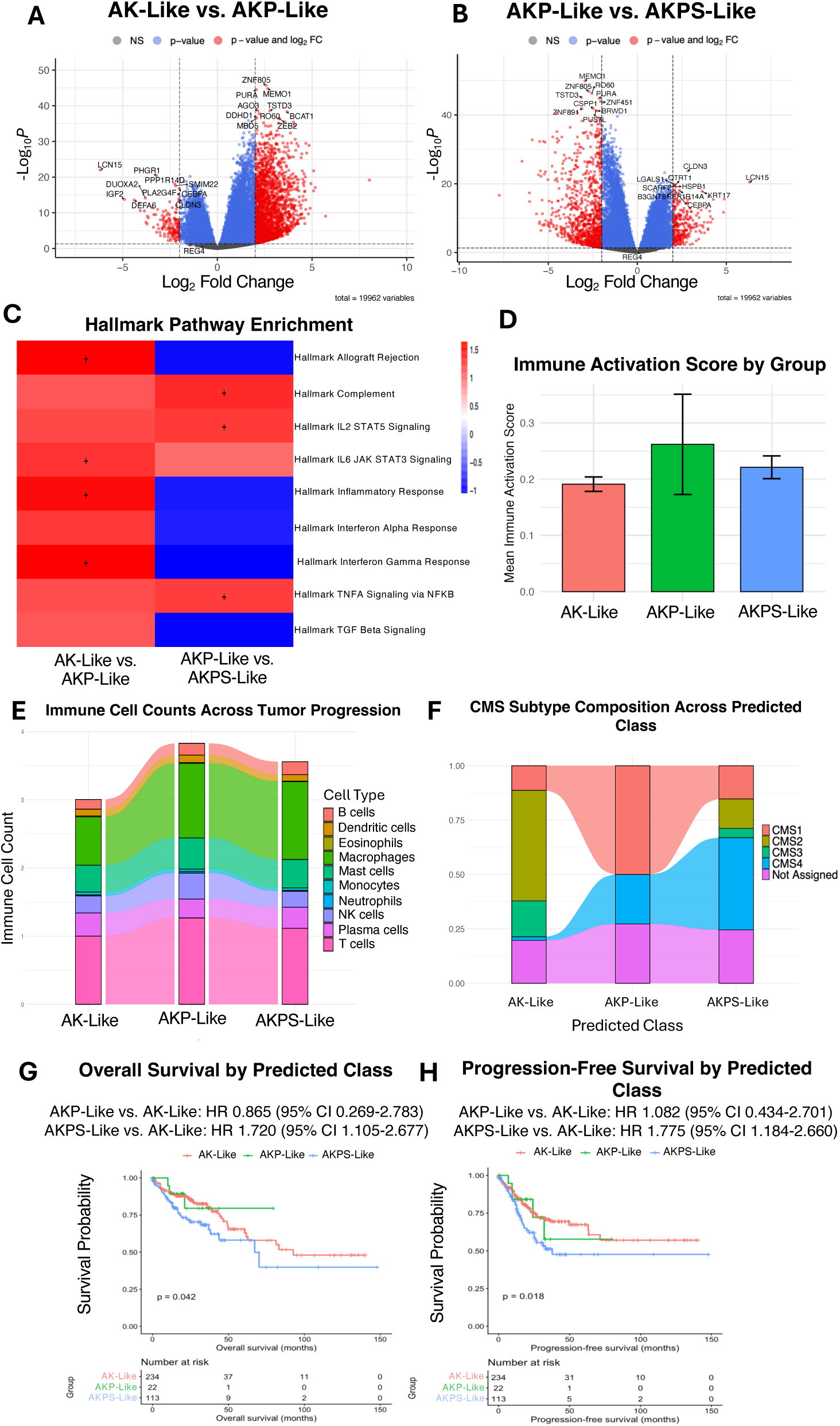
Transcriptomic and Gene Expression Changes Across Human Tumor Stages Defined by Cross-Species Classification. **(A–B)** Volcano plots showing differentially expressed protein-coding genes for AK-like vs. AKP-like (A) and AKP-like vs. AKPS-like (B) tumor groups, with labels assigned by cross-species classification. Genes meeting thresholds for both adjusted P value (< 0.05) and absolute log2 fold change (> 2) are shown in red; genes significant only by adjusted P value are shown in blue; nonsignificant genes are shown in gray. P values were calculated using the Wald test. **(C)** Heat map of NES for selected inflammatory Hallmark pathways identified by pre-ranked GSEA across AK-like vs. AKP-like and AKP-like vs. AKPS-like comparisons. A positive NES indicates enrichment in the later stage of each comparison. Significance was assessed using a permutation-based Kolmogorov–Smirnov statistic with gene set permutations, with false discovery rate (FDR) correction applied. **(D)** Bar plot showing mean immune activation scores across AK-like, AKP-like, and AKPS-like tumor groups, with error bars indicating variability within each group. **(E)** Alluvial plot depicting relative changes in immune cell subtype proportions across sequential human tumor stages defined by cross-species classification. **(F)** Alluvial plot showing changes in the proportion of Consensus Molecular Subtype (CMS) classes across the same sequential tumor stages, based on predicted CMS labels. **(G–H)** Kaplan–Meier curves showing overall survival (G) and progression-free survival (H) for AK-like, AKP-like, and AKPS-like tumor groups. Hazard ratios are shown relative to the AK-like group, and P values were calculated using univariate Cox proportional hazards models. **Biological and technical replicates:** Sample counts were AK-like (n = 248), AKP-like (n = 22), AKPS-like (n = 118), and unclassified (n = 72). Of 460 tumors analyzed, 458 represented unique patients; two patients contributed more than one biologically distinct tumor specimen (e.g., primary and metastatic).

These pathway-level dynamics were reflected in overall immune abundance. A composite immune activity score, derived from the summed contributions of multiple immune cell populations including CD8⁺ T cells, activated CD4 memory T cells, NK cells, macrophages, plasma cells, regulatory T cells, and neutrophils, showed significant elevation in AKP-like tumors relative to AK-like tumors, followed by a marked reduction in AKPS-like tumors (**Figure 8D**). However, these trends did not reach statistical significance.

Immune subtype analysis further clarified species-specific features of the conserved immune trajectory. Alluvial mapping showed that the early immune influx in human tumors with the loss of *TP53* was driven predominantly by macrophages, in contrast to the NK and T-cell-dominated influx observed in the mouse model (**Figure 8E**). Although the cellular drivers differed, both species exhibited a coordinated increase in innate and adaptive immune infiltration during the early transition, followed by broad immune suppression as they progressed to the AKPS-like state.

To determine whether the AK, AKP, and AKPS transcriptional states correspond to established biological subtypes of human CRC, we examined the distribution of CMS types (CMS1–CMS4) across the three groups (**Figure 8F**). CMS composition shifted in a highly structured, biologically coherent manner across states. The CMS overlay confirms the biological identity of the AK, AKP, and AKPS classes. AK aligns closely with CMS2/CMS3 and represents a stable epithelial baseline state. AKP is aligned with CMS1, revealing an intermediate immune-activated phenotype that bridges the epithelial-mesenchymal continuum. AKPS is almost entirely CMS4, consistent with an invasive, stromal-rich, late-stage phenotype.

To determine whether the conserved transcriptional and immune states identified above have clinical consequences in human CRC, we evaluated overall survival (OS) and progression-free survival (PFS) across AK-like, AKP-like, and AKPS-like tumors (**Figure 8G-H**). Kaplan-Meier analyses revealed that AKPS and AKP groups yielded worse OS or PFS in reference to AK tumors. Univariate Cox proportional hazards ratios (HR) demonstrated that tumors with the AKPS-like state exhibited a significant increase in risk, suggesting that this late-stage pathway disruption confers a marked survival disadvantage **(Figure 8G-H)**. In multivariable Cox models adjusted for American Joint Committee on Cancer (AJCC) stage and T/N/M classifications, the AKPS-like group remained statistically significant in decreased PFS (HR=1.58, 95CI=1.03-2.42, p=0.036) (**Supplementary Table 2-3**). This indicated that the AKPS-like transcriptional state provides independent prognostic value beyond N and M annotations, which were expectedly significant as these are clinical indicators of more advanced stages. In our multivariate analysis, the AK-like and AKP-like tumor groups no longer showed significant differences in PFS or OS, potentially due to the small sample sizes (**Supplementary Table 2-3**). Altogether, the transcriptional states, especially the AKPS-like group, capture prognostic information not explained by existing clinical classification schemes.

## DISCUSSION

Here, we define a temporally ordered transcriptional, immune, and stromal landscape accompanying the sequential acquisition of *APC, KRAS, TP53, and SMAD4* mutations in colorectal cancer (40). By reconstructing this canonical genetic trajectory in a controlled orthotopic organoid system, we uncover discrete, stage-specific biological state transitions that are not resolvable from cross-sectional human tumor datasets. CRC progression in this model is not characterized by cumulative amplification of early alterations, but instead by successive activation of distinct transcriptional programs accompanied by coordinated remodeling of the tumor microenvironment. These findings redefine CRC as a dynamic, co-evolving tumor–immune–stromal system rather than a static collection of genotypically defined entities. Consistent with this framework, human tumors reassigned into AK-like, AKP-like, and AKPS-like transcriptional states show biologically ordered and clinically significant differences in outcomes. Decreased progression-free survival was seen in the AKPS-like group, directly linking late-stage immune and stromal reprogramming to adverse clinical outcomes.

Analysis of large human transcriptomic cohorts, including TCGA (41), reveals substantial heterogeneity among tumors harboring identical *APC, KRAS, TP53, and SMAD4* alterations, with AK, AKP, and AKPS groups lacking clear transcriptional or immune demarcation. This unsurprisingly reflects fundamental limitations of human datasets, which capture tumors at a single time point and conflate mutation-driven effects with inter-patient and microenvironmental variability (42). Our organoid model resolves temporal ordering and controls for these environmental factors, therefore revealing insights that would be impossible to discern solely from human datasets.

Tumors derived from the organoid model progress through well-defined transcriptional states, consistent with a punctuated model of CRC evolution. Each mutational step induces a distinct reprogramming event with minimal overlap between stages, extending the Vogelstein framework (40) by integrating microenvironmental adaptation and phenotypic plasticity as core components of tumor progression. These transitions highlight that genetic alterations act not in isolation, but through emergent interactions with the surrounding immune and stromal compartments.

A key insight from this work is the identification of a temporally structured biphasic immune program coupled to EMT dynamics. Loss of TP53 in the APC-KRAS background induces a pronounced inflammatory state marked by interferon signaling and influx of effector immune populations, consistent with chromosomal instability and cGAS-STING activation (43). This immune activation coincides with early EMT induction, generating an inflamed yet plastic tumor state that may initially constrain tumor growth while imposing selective pressure for immune evasion (44).

Subsequent loss of SMAD4 precipitates a transition to an immunosuppressed, stromal-dominant state characterized by reduced lymphoid infiltration, expansion of fibroblast programs, contraction of EMT signatures, and disruption of TGF-β signaling (45). This shift promotes immune exclusion through myeloid skewing and extracellular matrix remodeling, culminating in immune-cold metastatic niches (46). Together, these findings define a conserved immune trajectory in CRC, in which early immunogenicity gives way to progressive immune dampening during advanced tumorigenesis.

The observed biphasic EMT program provides mechanistic insight into metastatic competence. Early EMT activation followed by EMT attenuation at metastatic sites supports a plasticity-based model in which transient EMT, rather than stable mesenchymal conversion, facilitates dissemination (47). Re-acquisition of epithelial features at metastatic sites is consistent with efficient colonization and aligns with emerging evidence from organoid and single-cell studies (48). Importantly, this EMT cycling integrates with immune remodeling, suggesting that phenotypic plasticity and immune state transitions are coupled processes during CRC evolution.

Our findings also reframe the biological interpretation of Consensus Molecular Subtypes (CMS). Rather than representing fixed tumor classes, CMS states emerge dynamically along the progression trajectory, with transitions from epithelial (CMS2) to immune-enriched (CMS1) to stromal-dominant (CMS4) phenotypes (3). In this context, CMS1 reflects a transient immune-activated state arising from coordinated tumor–immune interactions rather than a static identity defined by genomic alterations. This dynamic behavior explains mixed CMS assignments in patient samples and intra-tumoral CMS heterogeneity and highlights a limitation of considering CMS classifications alone when examining clinical outcomes (4).

Importantly, projecting mouse-derived progression modules onto human CRC uncovers conserved biological states that are not captured by mutational stratification alone. Human tumors align along the same immune, EMT, and stromal trajectories observed in the organoid system, including early immune activation and late immune suppression. While the dominant immune cell populations differ between species, the underlying evolutionary logic is preserved (bioRxiv 2022.10.02.508492), supporting progression-based classification as a robust framework for resolving tumor heterogeneity across systems. Notably, the AKPS progression state confers prognostic information beyond AJCC stage, underscoring the clinical relevance of progression-defined transcriptional identity.

Our insights have direct therapeutic implications. The immune-activated AKP/CMS1-like state represents a transient window of vulnerability during which tumors may be particularly responsive to immune engagement strategies, including innate immune activation or early checkpoint modulation. In contrast, the stromal-rich, TGFβ-dominated AKPS/CMS4 state suggests a need for microenvironment-targeted interventions to reverse immune exclusion. The dynamic nature of these states argues against fixed subtype-based treatment paradigms and highlights the importance of aligning therapy with the tumor’s current evolutionary state. The adverse clinical outcomes associated with the AKPS-like state underscore the urgency of targeting late-stage stromal and immune-suppressive programs, which appear to dominate disease progression once SMAD4-dependent regulation is lost.

Several limitations should be acknowledged. While the model captures a canonical genetic trajectory, it does not encompass the full mutational, metabolic, microbial, or host diversity of human CRC. Species-specific immune differences necessitate functional validation in human systems. Future studies integrating spatial transcriptomics, lineage tracing, and longitudinal sampling of early human lesions will be essential to validate these state transitions in patients.

## CONCLUSIONS

In conclusion, our study establishes a progression-resolved framework for CRC evolution (49), integrating sequential driver mutations with dynamic transcriptional reprogramming, immune state transitions, EMT-MET plasticity, stromal remodeling, and evolving CMS types. By shifting from static genotypes to emergent biological states, our data clarified human CRC heterogeneity, aligned preclinical organoid systems with clinical relevance, and uncovered temporally restricted therapeutic windows, such as exploiting early immune activation in AKP/CMS1-like tumors with checkpoint inhibitors or reversing late suppression in AKPS/CMS4-like cases with stromal disruptors. The strong concordance between progression-defined states and patient survival reinforces the clinical relevance of this framework and supports progression-based classification as a foundation for prognostic stratification and state-matched therapeutic intervention in CRC. Leveraging organoids as a reductionist platform to recapitulate stepwise progression surpasses limitations of heterogeneous human CRC datasets and conventional mouse models. At the same time, our bioinformatics approach, deriving progression modules and projecting them onto human samples via cross-species classification, validates conserved trajectories, including biphasic immunity and species-specific immune effectors. This scalable pipeline redefines mouse-human model evaluation across cancers.

## LIST OF ABBREVIATIONS

AK: tumors with APC/KRAS alterations
AKP: tumors with APC/KRAS/TP53 alterations
AKPS: tumors with APC/KRAS/TP53/SMAD4 alterations
CMS: consensus molecular subtype
CRC: colorectal cancer
EMT: epithelial-mesenchymal transition
FPKM: fragments per kilobase per million
GDC: Genomic Data Commons
GSEA: gene set enrichment analysis
H&E: Hematoxylin and Eosin
HR: hazards ratio
MGI: Mouse Gene Identifier
MmCMS: mouse-adapted CMS classifier
MSI: microsatelite instability
NCI: National Cancer Institute
NES: normalized enrichment score
OS: overall survival
PBS: phosphate buffered saline
PFS: progression free survival
TCGA: The Cancer Genome Atlas
TCGA-COAD: The Cancer Genome Atlas Colorectal Adenocarcinoma
TME: tumor microenvironment
TPM: transcripts per million
UMAP: Uniform Manifold Approximation and Projection
VST: variance stabilizing transformation

## DECLARATIONS

### Ethics approval and consent to participate

All animal studies proposed in this project were conducted in strict accordance with institutional and federal guidelines for the ethical use of vertebrate animals in research. All experimental protocols involving mice have been reviewed and approved by the Institutional Animal Care and Use Committee (IACUC) at the University of Minnesota.

### Consent for publication

Not applicable

### Availability of data and materials

The human colorectal cancer transcriptomic and associated clinical datasets analyzed during the current study are available from The Cancer Genome Atlas Colon Adenocarcinoma (TCGA-COAD) repository via the NCI Genomic Data Commons (GDC) data portal ^22^, https://portal.gdc.cancer.gov/analysis_page?app=Projects. The mouse RNA-sequencing datasets generated during the current study, including raw FASTQ files and processed expression matrices, are available in the Gene Expression Omnibus (GEO) repository under accession number GSE316489, https://www.ncbi.nlm.nih.gov/gds/. All other data generated and/or analyzed during this study are available from the corresponding author upon reasonable request.

The custom R scripts used in this study are publicly available. The archived version of the code associated with this manuscript is available at https://doi.org/10.5281/zenodo.18462555. The most recent version of the code is maintained at https://github.com/KhalidIshani/CRC_project.git.

### Competing Interests

EB consults for Tempus AI, Astrin Biosciences, and EMRGNSE. KS consults for EMRGNSE, LLC. JH consults for Tempus AI and Astrin Biosciences and is a co-founder of EMRGNSE, LLC. All other authors report no disclosures.

### Funding

Research funds from the Minnesota Colorectal Cancer Fund and Mezin Koat Colorectal Cancer Research Fund supported this study. The Minnesota Colorectal Cancer Research Foundation supported TJG and DW graduate fellowships.

### Author Contributions

KI performed the statistical analysis of the human/mouse data and played a major role in writing the manuscript. DM assisted with writing & editing the manuscript. AA helped with data preparation and conceptualization of the project idea. TG helped conceptualize the research idea and assisted with writing/editing of the manuscript. ZY performed the survival analysis and created figure 8. APG helped organize the project, assisted in figure preparation, and contributed to the writing/editing of the manuscript. EB assisted with data analysis and helped write/edit the manuscript. KS assisted with the survival analysis, writing the manuscript, and brainstorming statistical methods. PG reviewed and edited the manuscript. JH played a major role in the conceptualization of the research idea, project administration, and writing/editing the manuscript. SS developed the organoid model, helped organize the project, and played a major role in writing/editing the manuscript.

## Acknowledgements

We thank the Masonic Cancer Center, the Clinical and Translational Sciences Institute, and the Department of Surgery for their research support. We acknowledge the use of generative artificial intelligence tools to improve the clarity and readability of portions of the manuscript and to provide guidance during the development of R code for statistical analyses. All analyses, code implementation, data interpretation, and final decisions regarding content were performed by the authors, who take full responsibility for the accuracy, integrity, and originality of the work.

## Notes

https://portal.gdc.cancer.gov/analysis_page?app=Projects

https://www.ncbi.nlm.nih.gov/gds/

https://doi.org/10.5281/zenodo.18462555

https://github.com/KhalidIshani/CRC_project.git

