## Supplemental Methods for "Discrete Transcriptional States Define Biphasic Immune Response and Dynamic CMS Transitions in Colorectal Cancer"

**Mouse Organoid Culture**

Normal colon crypts were harvested from C57BL/6 WT mice, as previously described [O’Rourke, Kevin P., et al. “Isolation, culture, and maintenance of mouse intestinal stem cells.” *Bio-protocol* 6.4 (2016): e1733-e1733]. Briefly, colon tissue was isolated, longitudinally cut, and washed with cold PBS until the supernatant was clear of debris and fecal matter. The tissue was then washed with 5 mM EDTA in PBS to loosen the crypts. To further loosen and isolate the crypts, the colon tissue was vigorously pipetted up and down multiple times in cold PBS. Three fractions of the supernatant were collected and observed under a microscope. The fraction containing the most intact crypts was pelleted and resuspended in 30µL Matrigel (BD Biosciences; San Jose, CA, USA) and plated as a dome in a 12-well Tissue Culture-treated CytoOne plate (USA Scientific; Ocala, FL, USA). The plate was incubated at 37 °C for 10-15 minutes to allow the Matrigel to solidify. The *AK* (sh*Apc* and *Kras^G12D^* ) organoids were kindly provided by Dr. Khashayarsha Khazaie. AKP (sh*Apc*, *Kras^G12D^*, and *Trp53 ^KO^*) organoids were generated by introducing an additional *P53* knockout mutation in the AK organoids. The AKPS (sh*Apc*, *Kras ^G12D^*, *Trp53 ^KO^*, and *Smad4^KO^*) organoids were kindly provided by Dr. Peter Westcott [Westcott, Peter MK, et al. "Low neoantigen expression and poor T-cell priming underlie early immune escape in colorectal cancer." *Nature cancer* 2.10 (2021): 1071-1085]. The AK, AKP, and AKPS organoids were resuspended in 30µL Matrigel (BD Biosciences; San Jose, CA, USA) and plated as a dome in a 12-well Tissue Culture-treated CytoOne plate (USA Scientific; Ocala, FL, USA). The plate was incubated at 37 °C for 10-15 minutes to allow the Matrigel to solidify.

Normal colon and AK organoids were cultured in complete medium. The complete medium contain 1:1 ratio of advanced DMEMF/12 (AdDMEM, Thermo Fisher Scientific, Waltham, MA, USA) and conditioned L-WRN cell supernatant, containing 1X penicillin/streptomycin (Thermo Fisher Scientific, Waltham, MA, USA), 1 mM N-acetylcysteine (Sigma-Aldrich), 10 mM HEPES (Thermo Fisher Scientific, Waltham, MA, USA), 1X B27 (Life Technologies, Carlsbad, CA), 1X N2 (Invitrogen, Waltham, MA, USA), 1 mM N-acetylcysteine (Sigma-Aldrich, Burlington, MA, USA), 1X Glutamax (Thermo Fisher Scientific, Waltham, MA, USA), 100 μg/mL Primocin (Invivogen; San Diego, CA, USA), 10 μM SB202190 (Sigma-Aldrich, Burlington, MA, USA) 10 μM Y-27632 (EMD Millipore; Burlington, MA, USA), 50 ng/mL human EGF (Invitrogen, Waltham, MA, USA), and 10 mM Nicotinamide (Sigma-Aldrich; Burlington, MA, USA). The AKP organoids were cultured in the same complete media as normal colon organoids with the addition of 13μM of Nutlin-3a (Cayman Chemical; Ann Arbor, MI, USA). The AKPS organoids were cultured in minimal media (AdDMEM F/12 supplemented with 1X B27). When the organoids reached confluency, they were treated with TrypLE Express at 37 °C for 10 minutes, neutralized with complete media/minimal media, and plated as domes in a new 12-well plate. The medium was replenished every 2-3 days.

**Colonoscopy-guided organoid injection**

Colonoscopy-guided orthotopic injection of organoids was performed, as previously described [Roper, Jatin, et al. "Colonoscopy-based colorectal cancer modeling in mice with CRISPR–Cas9 genome editing and organoid transplantation." Nature protocols 13.2 (2018): 217-234]. Two days before the injection, organoids were broken into small aggregates in TrypLE Express at 37 °C for 5 minutes by pipetting up and down. Organoids were washed in conditioned L-WRN cell supernatant (L-WRN conditioned media) and centrifuged at 300g for 5 minutes. The pelleted organoids were washed in PBS, centrifuged at 300g for 5 minutes, and then resuspended in minimal media with 10% Matrigel at ~50 organoids/μL. Mice were anesthetized using isoflurane. Organoids were orthotopically injected via a Hamilton syringe through the working channel of the colonoscope. The tip of the injection needle was inserted into the mucosa of the colon at ~30° angle relative to the colon wall. Successful injection of organoids should result in a small bubble form visible within the mucosa.
